# Pan-Cancer Genomic Scars of Alternative End Joining and Single-Strand Annealing

**DOI:** 10.64898/2026.05.22.727283

**Authors:** Ashini Modi, Alessandro Zito, Giovanni Parmigiani

## Abstract

DNA double-strand breaks (DSBs) are among the most cytotoxic lesions encountered by cells and represent a major source of genome instability in cancer. The preferred pathway to repair resected DSBs is high-fidelity homologous recombination (HR), but error-prone annealing-dependent pathways such as single-strand annealing (SSA) and alternative end joining (Alt-EJ) generally act as backup repair mechanisms in HR-deficient contexts. However, the extent to which these pathways are engaged across tumor types and how their activity is coupled with HR deficiency remains unclear. In this work, we systematically characterize scars from SSA and Alt-EJ across 2,157 whole-genome-sequenced tumors spanning 17 cancer types, identifying 832 SSA-like and 37,359 Alt-EJ-like deletions in total. We find that Alt-EJ is the predominant backup repair pathway in HR-deficient tumors compared with SSA; however, prostate adenocarcinoma and hepatocellular carcinoma exhibit elevated SSA-like deletion burdens despite low HR-deficiency scores. Moreover, our genome-wide analysis reveals that SSA-like deletions preferentially occur in SINE-rich regions and exhibit pronounced enrichment near transcription start sites in HR-proficient lymphoid lineage tumors. Our results show that SSA- and Alt-EJ-associated genomic scars are not confined to HR-deficient tumors, but are shaped by local genomic and transcriptional context, capturing distinct dimensions of DSB repair activity beyond HR deficiency alone.

## Introduction

DNA double-strand breaks (DSBs) are highly mutagenic lesions that, if repaired incorrectly, can lead to deletions, duplications, focal amplifications, or chromosomal translocations. These aberrations strongly contribute to carcinogenesis by deleting tumor suppressor genes, amplifying oncogenes, or repositioning regulatory elements to induce inappropriate gene expression (Lengauer et al. 1998; Mladenov et al. 2016; Mills et al. 2003; Wang et al. 2020). Therefore, characterizing the drivers and frequency of mutagenic DSB repair is central to understanding the origins of genomic instability in cancer.

Cells employ multiple mechanistically distinct pathways to repair DSBs, each differing in fidelity, timing, and mutagenic potential. Canonical non-homologous end joining (c-NHEJ) directly ligates DNA ends with minimal sequence homology, while homologous recombination (HR) uses a homologous template to restore sequence information and is largely error-free (Scully et al. 2019; Ceccaldi et al. 2016; Pannunzio et al. 2018; Jasin and Rothstein 2013). Resected DSBs can also be repaired through annealing-dependent pathways, including single-strand annealing (SSA) and alternative end joining (Alt-EJ) (Ceccaldi et al. 2016). SSA, mediated by Rad52, anneals homologous repeat sequences flanking a DSB, resulting in deletion of one repeat and the intervening sequence, often spanning kilobases of DNA (Bhargava et al. 2016; Blasiak 2021). In contrast, Alt-EJ, also known as microhomology-mediated end joining (MMEJ), often mediated by POLQ, uses short microhomologies to rejoin DNA ends and is associated with small deletions, insertions, and complex junctional architectures. Both SSA and Alt-EJ are intrinsically mutagenic, leading to irreversible loss of genetic information (Sallmyr and Tomkinson 2018).

SSA and Alt-EJ engage as backup pathways upon loss of HR or c-NHEJ. In HR-deficient cells, Alt-EJ activity is associated with increased POLQ expression and enrichment of microhomology-mediated insertions and deletions, including mutational signatures such as ID6 (Ceccaldi et al. 2015; Bazan Russo et al. 2024; Alexandrov et al. 2018). In parallel, impairment of HR, including loss of RAD51 or BRCA1/2 function, has been shown to promote reliance on SSA (Stark et al. 2004; Blasiak 2021; Setton et al. 2023) while inhibition of the SSA mediator RAD52 selectively reduces viability of HR-deficient cells (Lok et al. 2013). However, accumulating evidence suggests that annealing-dependent repair pathways are not merely emergency substitutes for HR, but are also engaged to varying degrees in HR-proficient contexts (Deriano and Roth 2013; Mateos-Gomez et al. 2015; Blasiak 2021). Indeed, the choice of DSB repair pathway is governed by a complex interplay among end resection, chromatin state, transcriptional activity, and local sequence architecture. For instance, when DSB occurs in the S phase in the absence of sister chromatids, it may be repaired by SSA or Alt-EJ due to a lack of a preferred template for HR (Johnson 2000). Furthermore, more closed chromatin impedes the building of a multi-protein complex required for HR at the damage site, while SSA proteins may be compatible with a less accessible chromatin (Blasiak 2021). As a result, error-prone repair pathways can be preferentially utilized in specific genomic or cellular environments, contributing to mutagenesis even when HR is functionally intact.

Despite growing recognition of the role of SSA and Alt-EJ in shaping cancer genomes, key questions remain unresolved. How frequently are these pathways used across tumor types? To what extent do SSA and Alt-EJ operate alongside HR rather than solely compensating for its loss? Addressing these questions is critical, as compensatory repair pathways can undermine therapies targeting DDR defects and influence both tumor evolution and treatment response. This paper aims to fill this gap. In particular, we systematically characterize the usage of SSA and Alt-EJ DSB repair pathways across 2,157 tumors from the International Cancer Genome Consortium (ICGC) database spanning 17 cancer types. By quantifying pathway-specific genomic scars and examining their relationship to HR proficiency, we aim to disentangle how SSA and Alt-EJ repair mechanisms contribute to cancer genome evolution and to clarify the role of DSB repair pathway choice in tumor etiology.

## Results

### Alt-EJ and SSA-like deletions increase with homologous recombination deficiency

To systematically characterize deletion-associated repair processes, we developed an algorithm (Methods) to detect deletions flanked by either microhomology or extended homeology. A summary of the classification for SSA and Alt-EJ is reported in Fig. 1. We classify deletions as consistent with alternative end-joining (Alt-EJ) or single-strand annealing (SSA) based on sequence features at the breakpoints. Deletions were classified as Alt-EJ-like if they exhibited microhomology lengths between 2 and 25 bp and deletion lengths greater than 5 bp. As for SSA-like events, we searched for direct repeats flanking deletion breakpoints, requiring a minimum repeat length of 30 bp and at least 80% sequence identity.

**Fig. 1.**
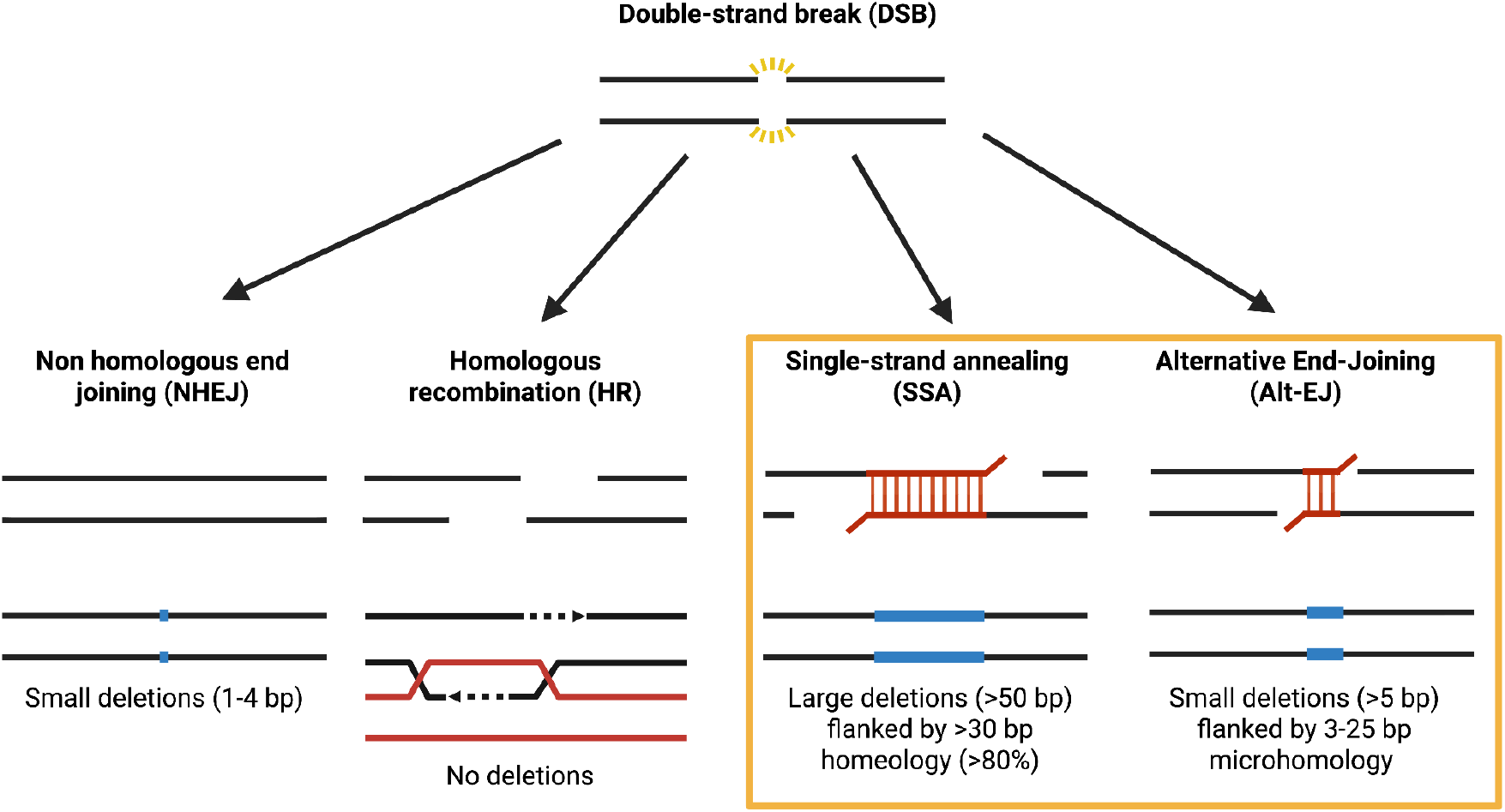
Major pathways for double-strand break (DSB) repair and their characteristic deletion signatures. Following a DSB, cells repair DNA through four primary pathways. Non-homologous end joining (NHEJ) directly ligates broken ends with minimal processing, typically generating small insertions or deletions (1-4 bp). Homologous recombination (HR) uses an intact homologous template to restore the sequence accurately, resulting in no deletions. Single-strand annealing (SSA) utilizes long homologous repeats; in our algorithm, we define repeats as >30 bp with ≥80% sequence identity, leading to large deletions (>50 bp) between repeats. Alternative end-joining (Alt-EJ), also known as microhomology-mediated end joining, relies on short microhomologies (defined here as 3-25 bp) and produces intermediate-sized deletions (>5 bp).

With this classification scheme, we identified 832 SSA-like and 37,359 Alt-EJ-like deletions. Across all tumors, Alt-EJ-like deletions exhibited a peak length of approximately 5 bp, whereas SSA-like deletions showed a peak at approximately 5 kbp, with substantially broader size distributions. Notably, SSA-like events displayed marked heterogeneity across tumors, with homeology tracts extending up to 392 bp (Supplementary Fig. 1). Quantitatively, 20.67% of deletions between 5-99 bp satisfied microhomology criteria and were classified as Alt-EJ-like, whereas only 3.76% of deletions between 66 bp and 707,723 bp satisfied homeology criteria and were classified as SSA-like, indicating that SSA events are substantially rarer.

To relate these patterns to homologous recombination deficiency (HRD), we compute HRD scores using telomeric allelic imbalance, loss of heterozygosity, and large-scale state transitions. Among deletions <100 bp, events of 6-20 bp (Fig. 2D1) with 2-5 bp microhomology (Fig. 2D2) showed the strongest positive correlation with HRD. In contrast, among deletions >100 bp, events in the 1-10 kbp range (Fig. 2E1) with 31-100 bp homeology (Fig. 2E2) were most strongly associated with HRD. These patterns are consistent with distinct mechanistic regimes underlying Alt-EJ and SSA repair, supporting the validity of our classification thresholds.

**Fig. 2.**
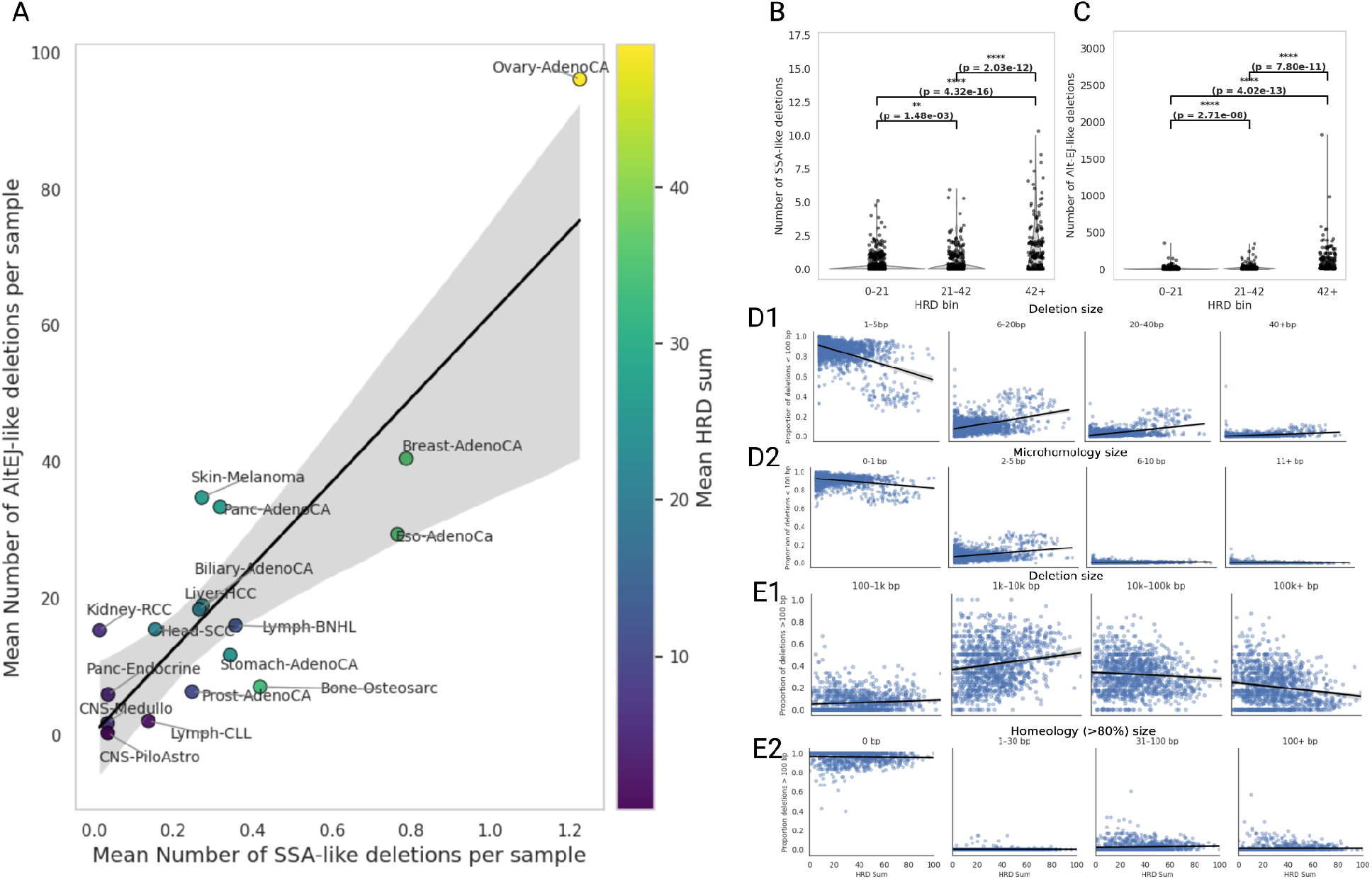
SSA and Alt-EJ deletions scale with homologous recombination deficiency. **a**, Mean number of Alt-EJ–like deletions plotted against mean number of SSA-like deletions per sample across cancer types, colored by mean HRD score. The line indicates the line of best fit, with the shaded region representing the 95% confidence interval. **b,c**, Distribution of SSA-like **(b)** and Alt-EJ–like **(c)** deletions across HRD bins (0-21, 21-42, >42). Points represent individual tumors. P-values are calculated using pairwise Welch’s tests between HRD groups. **d,e**, Relationship between deletion features and HRD score. **(d1)** Proportion of deletions <100 bp stratified by size (1-5 bp, 6-20 bp, 20-40 bp, >40 bp) as a function of HRD score. **(d2)** Proportion of deletions <100 bp stratified by microhomology length (0-1 bp, 2-5 bp, 6-10 bp, ≥11 bp). **e1,e2**, Proportion of deletions >100 bp (**e1**) stratified by size (100-1 kb, 1-10 kb, 10-100 kb and >100 kb) and (**e2**) homology length (>80% identity; 0 bp, 1-30 bp, 31-100 bp and >100 bp) as a function of HRD score. Each point represents a tumor sample, and lines indicate linear fits.

Both Alt-EJ-like and SSA-like deletion burdens increased with HRD score, confirming that both backup repair pathways are increasingly relied upon in HR-deficient tumors. Ovarian adenocarcinoma exhibited the highest mean HRD score (∼49.1), alongside elevated mean Alt-EJ-like (∼96.1 per tumor) and SSA-like (∼1.2 per tumor) event counts (Fig. 2A). Consistent with this, tumors harboring BRCA2, but not BRCA1, loss-of-function mutations exhibited significantly higher Alt-EJ-and SSA-like deletion burdens compared to wild-type tumors, with a more pronounced increase than observed for BRCA1-deficient tumors (Fig. S2). These results are in line with prior work demonstrating that SSA and Alt-EJ are preferentially active in BRCA2-deficient cancers (Han et al. 2017; Tutt 2001; Stark et al. 2004; Blasiak 2021; Setton et al. 2023).

While a threshold of 42 is commonly used to define HR deficiency (Telli et al. 2023), tumors with intermediate HRD scores (21-42) displayed significantly elevated deletion burdens relative to HR-proficient tumors (0-21), yet lower than HR-deficient tumors (>42), supporting the interpretation of this range as a biologically relevant intermediate repair phenotype (Fig. 1B,C).

### Cancers stratify by SSA and Alt-EJ usage independent of HRD status

We next examined how the relationship between usage of SSA and Alt-EJ repair pathways and HRD varies across tumor types. In particular, both Alt-EJ-like and SSA-like deletion counts increased with HRD score for many cancer types, consistent again with increased reliance on backup repair pathways in HR-deficient genomes (Fig. 3A, B). An exception was observed in melanoma, which showed a lower median Alt-EJ burden in the HRD-high (42+) group compared to HR-proficient tumors (0-21) (Fig. 3B).

**Fig. 3.**
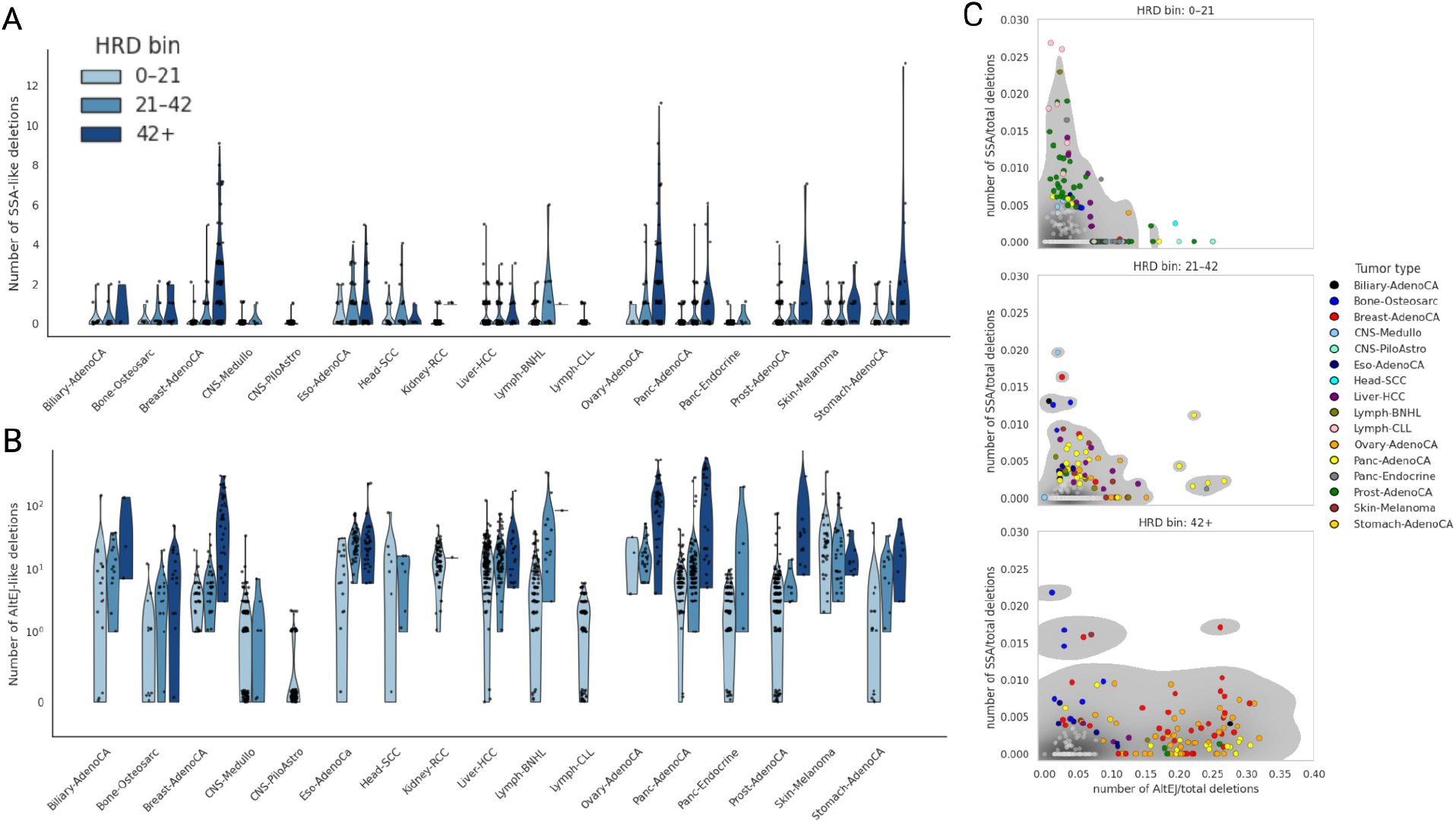
SSA and Alt-EJ usage across tumor types and HRD bins. **a**, Distribution of SSA-like deletions across tumor types stratified by HRD bin (0-21, 21-42, >42). **b**, Distribution of Alt-EJ-like deletions across tumor types stratified by HRD bin. Points represent individual tumors overlaid on violin plots showing the distribution within each cohort. **c**, Relationship between normalized SSA and Alt-EJ burden across HRD bins. Each point represents a tumor, colored by tumor type. Axes show the proportion of SSA-like deletions (y axis) and Alt-EJ–like deletions (x axis), normalized by the total number of deletions per tumor as a proxy for genomic instability. Grey density contours indicate the distribution of samples within each HRD bin, and colored points denote tumors outside the central high-density region (defined as the region containing 99% of the density mass).

Within HR-proficient tumors (HRD 0-21), most tumor types exhibited few cases with detectable SSA activity (75th percentile = 0 events); however, prostate adenocarcinoma and liver hepatocellular carcinoma showed increased fractions of tumors harbouring at least one SSA-like deletion (Fig. 3A). Both tumor types were significantly enriched relative to other cohorts (prostate: 15.5%, liver: 16.2% versus 8.6%; odds ratios = 1.96 and 2.07, respectively; Fisher’s exact test, *P* = 0.0064 and *P* = 0.0021), indicating that SSA-like repair is preferentially engaged in these tumor types even in the absence of HR deficiency.

To quantify Alt-EJ activity, we defined high Alt-EJ burden as the upper quartile (75th percentile: 10 events per tumor) and compared enrichment at this threshold. Melanoma, kidney, and liver tumors showed strong enrichment for high Alt-EJ burden (81.8%, 75.3%, and 68.4% of tumors, respectively; odds ratios = 15.0, 11.3, and 10.0; Fisher’s exact test, *P* < 10^−12^) (Fig. 3B).

SSA and Alt-EJ events were also normalized by the total number of deletions per tumor to account for variation in overall genomic instability (Fig. 3C). In HR-proficient tumors, the joint distribution of normalized values showed a pronounced tail of samples with elevated SSA relative to Alt-EJ. As HRD increases, this SSA-enriched tail diminishes, whereas a corresponding tail of Alt-EJ-enriched tumors expands, indicating a shift toward preferential Alt-EJ usage under HR-deficient conditions. This is in line with experimental evidence supporting a hierarchical organization of DSB repair pathways, in which Alt-EJ functions as a backup when HR and c-NHEJ are compromised (Iliakis et al. 2015). We identified samples outside the central distribution (top 1% of proportions), corresponding to SSA-enriched or Alt-EJ-enriched outliers. Prostate tumors were overrepresented among HR-proficient SSA-enriched outliers, whereas breast, ovarian, and pancreatic tumors were overrepresented among HR-deficient Alt-EJ-enriched outliers.

### Genomic hotspots of Alt-EJ and SSA-like deletions

We next examined the spatial distribution of these events across the genome to determine the presence of tumor-specific regions of enrichment. We identify two hotspots of Alt-EJ genomic scars (Fig. 4A). In lymphoid malignancies (Lymph-BNHL and Lymph-CLL), Alt-EJ events were strongly enriched at chromosome 14q32, a locus encompassing the immunoglobulin heavy chain (IGH) region. This enrichment is consistent with the known role of alternative end-joining in mediating illegitimate joining of DNA breaks during V(D)J recombination, generating oncogenic translocations such as t(14;18) (Marculescu et al. 2002). A second enrichment region was observed on chromosome 4q13 in liver tumors. This region overlaps a locus frequently affected by loss of heterozygosity in hepatocellular carcinoma (Niu et al. 2016), suggesting that Alt-EJ-mediated repair may provide a mechanistic basis for the recurrent deletions observed across 4q13.3-q35.2 and the associated loss of tumor suppressor genes.

**Fig. 4.**
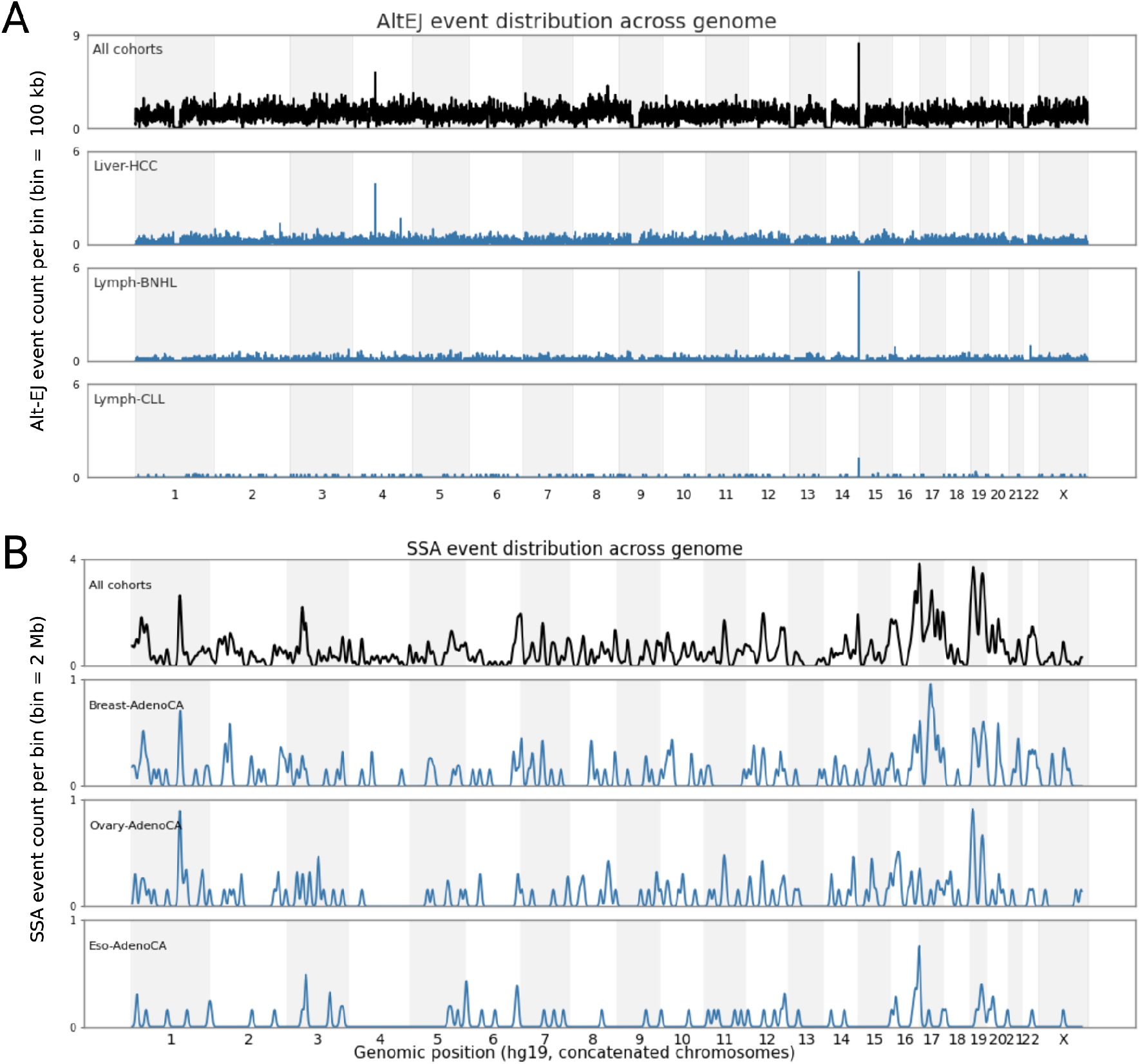
Genome-wide distribution of Alt-EJ and SSA events across tumor types. **a**, Genome-wide distribution of Alt-EJ–like deletions, with chromosomes plotted sequentially along a continuous hg19 coordinate axis. The top panel (black) shows all cohorts combined, and lower panels (blue) show representative tumor types (top, Liver-HCC; middle, Lymph-BNHL; bottom, Lymph-CLL). Event density is shown as Alt-EJ event counts per 100 kb bin. **b**, Genome-wide distribution of SSA-like deletions across the genome (hg19). The top panel (black) shows all cohorts combined, and lower panels (blue) show representative tumor types (top, Breast-AdenoCA; middle, Ovary-AdenoCA; bottom, Eso-AdenoCA). Event density is shown as SSA event counts per 2 Mb bin.

SSA breakpoint density was positively correlated with SINE repeat density (Spearman ρ = 0.30, *P* = 3.1 × 10^−15^). This is because chromosomal translocations occur at high frequency when a DSB lies between neighboring Alu repeats (Elliott et al. 2005; Morales et al. 2015). Moreover, enrichment of SSA-like deletions was observed on chromosome 16q22-q24 in both esophageal and breast tumors (Fig. 4B), and weakly in pancreatic and prostate tumors (Supplementary Fig. 3). In breast adenocarcinoma, further enrichment was observed at chromosome 17q21, encompassing key loci including BRCA1 (Reinholz et al. 2009), as well as at chromosome 1q21-q22 and 19q13. We also detected similar enrichment at chromosomes 1 and 19 in ovarian tumors (Fig. 4B). Genome-wide distributions of SSA and Alt-EJ-like deletions across all cancer types are shown in Supplementary Fig. 3.

### SSA-like deletions are transcriptionally associated in HR proficient tumors

DSBs are not uniformly distributed across the genome, but are enriched in regions of active transcription. We therefore investigated whether SSA- and Alt-EJ-like deletions show preferential localization relative to transcription start sites (TSS). Fig. 5 shows the density of distances between deletion breakpoints and the nearest TSS. SSA-like deletions in HR-proficient tumors (HRD 0-21) were strongly enriched at TSS, exhibiting a sharp peak centered at 0 kb that is not observed for larger deletions (>100 bp) or for deletions more generally (Fig. 5A, B). The TSS enrichment in HRD 0-21 is primarily driven by lymphoid malignancies (Supplementary Fig. 4A). This is consistent with prior experimental evidence that SSA may be essential in the pathogenesis of hematopoietic cancers (Wilch and Morton 2018). In contrast, in HR-deficient tumors, the TSS-centered enrichment was attenuated, consistent with more widespread usage of SSA under conditions of elevated DNA damage. Alt-EJ-like deletions also showed moderate enrichment at TSS across HRD groups, exceeding that observed for deletions <100 bp overall (Fig. 4C-D, Supplementary Fig. 4C).

**Fig. 5.**
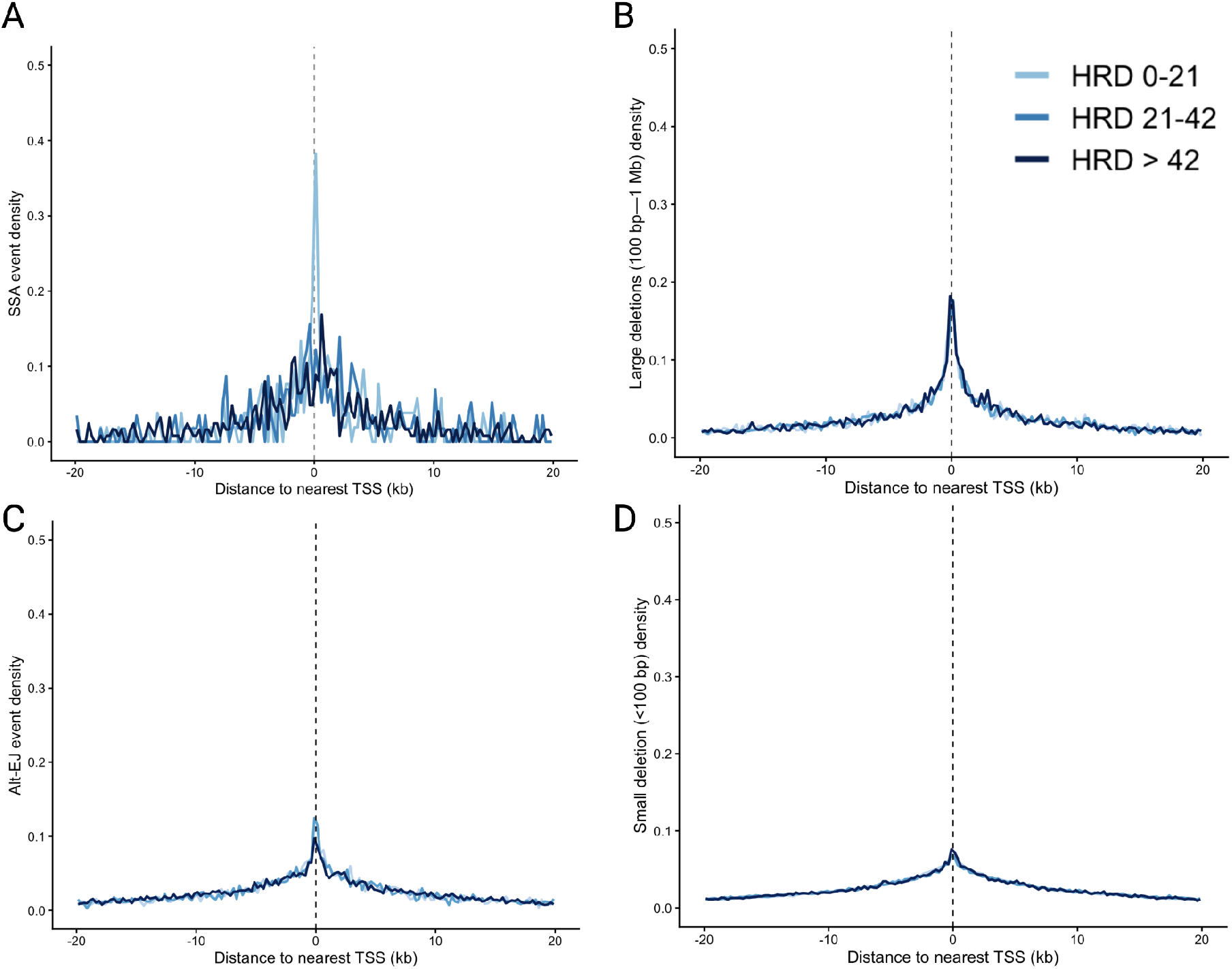
Distribution of deletion breakpoints relative to transcription start sites. **a**, Density of SSA-like deletion breakpoints as a function of distance between the nearest breakpoint and the nearest transcription start site (TSS), stratified by HRD bin (0-21, 21-42, >42). **b**, Density of large deletions (≥100 bp) relative to TSS across HRD bins. **c**, Density of Alt-EJ-like deletion breakpoints relative to TSS across HRD bins. **d**, Density of small deletions (<100 bp) relative to TSS across HRD bins. In all panels, distance is measured from the nearest breakpoint to the nearest TSS, with 0 kb indicating the TSS position (dashed line). Breakpoint distances were binned into 0.25-kb intervals within a ±20-kb window around TSSs. Curves represent normalized event densities within each HRD bin pooled across all tumors.

## Discussion

We systematically characterized scars of the SSA and Alt-EJ pathways across 2,157 whole-genome sequenced tumors spanning 17 cancer types. Our analysis identified 832 SSA-like and 37,359 Alt-EJ-like deletions. Our study provides the largest genome-wide quantification of SSA and Alt-EJ deletion burden across cancers using WGS, enabling direct comparison of pathway usage across varying degrees of HR-proficiency.

We observe a marked increase in Alt-EJ-associated events with increasing HRD score, whereas SSA increases more weakly. This suggests preferential engagement of Alt-EJ as a compensatory pathway in HR-deficient genomes. Such an increase parallels prior pan-cancer analyses showing that indel signature ID6, defined by microhomology-flanked deletions, co-occurs with SBS3 exposure and BRCA1/2-associated homologous recombination deficiency, implicating Alt-EJ as a dominant process in HRD tumors (Alexandrov et al. 2018). Microhomology has also been observed at breakpoint junctions of oncogenic chromosomal translocations in primary tumors, supporting a role for Alt-EJ in the formation of these rearrangements and, more broadly, in cancer pathogenesis (Rass et al. 2009; Zhang and Jasin 2011).

We further demonstrate that both SSA and Alt-EJ are utilized in HR-proficient contexts in a tumor-specific manner. Alt-EJ, while initially considered a backup pathway, has been shown to operate alongside canonical non-homologous end joining (c-NHEJ) and homologous recombination (HR), and may represent the dominant repair route for specific classes of DNA damage (Schrempf et al. 2021; Dueva and Iliakis 2013; Smolinska et al. 2022). Consistent with this, we observe elevated Alt-EJ activity in liver, kidney RCC, and melanoma despite low HRD scores (0-21). In contrast to Alt-EJ, whose role in cancer genomes is increasingly well established, the contribution of SSA has remained less clearly defined at the genome-wide level. In reporter assays, RAD52-dependent SSA is strongly upregulated upon RAD51 inhibition (Bennardo et al. 2008), and RAD52 loss is synthetically lethal with BRCA1/2 deficiency (Lok et al. 2013). More recently, scars of SSA repair have been observed in BRCA2-deficient tumors (Setton et al. 2023), however, their relevance across diverse tumor types, and particularly in HR-proficient cancer genomes, has not been systematically demonstrated. Our results extend these findings by demonstrating that SSA-associated deletions are not restricted to HRD tumors, but are also present across HR-proficient cancer genomes lacking canonical DDR defects. Given that SSA-mediated repair is intrinsically mutagenic and has been linked to repeat-mediated chromosomal rearrangements, these findings suggest that SSA may be a relevant contributor to cancer transformation independently of HR loss.

At a finer scale, SSA-like deletions are strongly enriched at transcription start sites (TSS) in HR-proficient tumors, with a sharp peak centered at 0 kb that is not observed for larger deletions or for deletions more broadly. This enrichment is driven by lymphoid malignancies, implicating SSA in the repair of transcription-associated DSBs in these tumors.

Current HRD classifiers rely heavily on copy-number alterations and mutational signatures, such as SBS3, yet these features do not fully capture the diversity of repair-pathway usage. Indeed, a substantial fraction of tumors exhibiting HRD-like genomic features lack canonical mutations in HR genes (McGrail et al. 2023). Our results suggest that incorporating SSA- and Alt-EJ-specific genomic scars may improve the resolution of these classifiers and better capture underlying repair phenotypes. Both SSA and Alt-EJ have been proposed as targets for synthetic lethality in HR-deficient cancers, with RAD52 inhibition impairing survival in BRCA1/2-deficient cells (Toma et al. 2019) and POLQ inhibition selectively targeting HRD tumors (Ronson et al. 2023). Our finding that these pathways are active even in HR-proficient tumors raises the possibility that they may support more precise stratification of repair phenotypes and inform the rational use of DNA-damaging therapies. More broadly, our results support a shift from viewing SSA and Alt-EJ solely as backup pathways to recognizing them as integral contributors to DSB repair and mutagenesis across diverse genomic contexts.

In this work, we inferred repair-pathway usage from sequence features rather than directly measuring it. In the future, it will be worth complementing our findings via orthogonal data, such as pathway-specific protein expression, including POLQ for Alt-EJ/MMEJ and RAD52 for SSA. Importantly, the boundary between Alt-EJ and SSA is not sharply defined biologically, and the exact repeat length that marks the transition between the two pathways is unclear. Experimental studies suggest that Alt-EJ typically utilizes microhomologies of ∼2-20 bp, whereas SSA involves longer homologous regions often exceeding ∼25 bp (Sallmyr and Tomkinson 2018). As a result, the repeat thresholds used here (>2 bp for Alt-EJ-like and ≥30 bp for SSA-like) are operational definitions chosen to reduce overlap between categories, though some ambiguity may remain for events near these boundaries.

## Methods

### Data

We analyzed a total of 2,157 tumors from the unrestricted access ICGC database spanning 17 cancer types, including breast, ovarian, pancreatic, prostate, stomach, and esophageal adenocarcinomas, pancreatic neuroendocrine cancer, squamous cell carcinoma, liver hepatocellular carcinoma, non-hodgkins lymphoma, chronic lymphocytic leukemia, melanoma, osteosarcoma, renal cell carcinoma, medulloblastoma, and pilocytic astrocytoma (Zhang et al. 2019).

Data was downloaded from University of California Santa Cruz Xena browser at https://xenabrowser.net/datapages/?cohort=PCAWG%20%28donor%20centric%29&removeHub=https%3A%2F%2Fxena.treehouse.gi.ucsc.edu%3A443 and is aligned to the GRCh37/hg19 reference genome.

### Classification of Alt-EJ-like Deletions

Alt-EJ, also known as microhomology-mediated end joining and Theta-mediated end joining, typically yields deletions with short microhomology tracts at the junction and deletion sizes beyond single-base indels (Setton et al. 2021). We therefore classified Alt-EJ-like deletions from indel callsets based on deletion size and junction microhomology length. For each indel record annotated as a deletion, we extracted a 50-bp upstream flank immediately upstream of the deletion start (hg19 positions -50 to -1 from the start of the deletion) and a 50-bp downstream flank immediately downstream of the deletion end (hg19 positions 1 to 50 from the deletion end).

Microhomology length was defined as the longest contiguous sequence that could plausibly anneal across the deletion junction. This consists of the maximum of the longest suffix of the deleted sequence matching the end of the upstream flank and the longest prefix of the deleted sequence matching the start of the downstream flank. For each deletion, candidate matches were searched by iteratively shortening the deleted sequence until a match to either flank was found, and the maximum match length across the upstream-anchored and downstream-anchored searches was recorded as the microhomology length. For indel deletions, we define length as the number of deleted reference bases, corresponding to the length of the reference allele for deletion calls.

A deletion was classified as Alt-EJ-like if it met both criteria of microhomology length between 3-25 bp and deletion length greater than 5 bp (Ceccaldi et al. 2016; Wyatt et al. 2016; Hussain et al. 2021; Seol et al. 2018). These thresholds were selected based on the ID6 mutational signature, thought to be the signature of Alt-EJ repair (Helleday et al. 2014), and to exclude blunt or near-blunt ligations as well as very short events consistent with c-NHEJ.

### Identification of SSA-like Deletions

SSA requires extensive resection and annealing between long homologous direct repeats, generating large deletions flanked by long, high-identity repeats. From indel callsets, candidate SSA substrates were restricted to deletion events with microhomology size greater than 30 bp (Setton et al. 2021; Seol et al. 2018; Ceccaldi et al. 2016). This minimum size threshold was selected to avoid overlap with the regime dominated by Al-EJ events. From the PCAWG consensus SV table, we restricted to events annotated as deletions.

For each SV deletion candidate, we extracted hg19 flanking sequences on both sides of the breakpoint using a flank size of 500 bp. Specifically, we extracted an upstream flank from positions Start™N to Start™1 and a downstream flank from positions End+1 to End+N with N = 500 bp. To classify an event as SSA, we searched for the longest shared sequence between upstream and downstream flanks that met both a minimum repeat length of at least 30 bp (Ceccaldi et al. 2016) and a minimum percent identity of at least 80% (Blasiak 2021; Setton et al. 2023). Percent identity was computed as the fraction of matching aligned bases over the candidate repeat length, and the longest repeat satisfying these thresholds was recorded for each deletion. Deletions meeting both thresholds were labeled SSA candidates. We performed local pairwise alignment of upstream versus downstream flanks using Biostrings pairwiseAlignment in local mode, and extracted aligned length and percent identity from the highest-scoring homologous segment.

### HRD Status calculation

Homologous recombination deficiency was quantified using the ‘scarHRD’ R package (Sztupinszki et al. 2018; Davies et al. 2017). Allele-specific copy-number, tumor purity, and ploidy estimates were obtained from the donor-centric copy-number data provided by the Pan-Cancer Analysis of Whole Genomes (PCAWG) data portal and were aligned to the GRCh37/hg19 reference genome. scarHRD was applied to allele-specific copy-number segmentations to compute three established genomic scar metrics: loss of heterozygosity (LOH), large-scale transitions (LST), and the number of telomeric allelic imbalances (TAI). For each tumor sample, an aggregate HRD score (HRD-sum) was defined as the sum of these three components. We restricted the analysis to tumors with HRD scores between 0 and 100 to exclude extreme outliers, which may result from sample processing rather than biological variation.

### Mapping Breakpoints to Transcription Start Sites

To assess the spatial relationship between DSB repair-associated SVs and transcription, we computed the genomic distance between SV breakpoints and the nearest transcription start site (TSS). A TSS was defined using the hg19 knownGene annotation from the UCSC Genome Browser (TxDb.Hsapiens.UCSC.hg19.knownGene). For each annotated transcript, the TSS was defined as the strand-aware 5’ end of the transcript and represented as a single-base genomic coordinate. For each event, two breakpoint coordinates were generated corresponding to the left and right boundaries of the deletion. For each breakpoint, the nearest TSS was identified using a nearest-neighbor search implemented with GenomicRanges. The distance between the closest breakpoint and its nearest TSS was calculated as a signed genomic distance, defined as the breakpoint coordinate minus the TSS coordinate. Thus, negative values indicate breakpoints upstream of the TSS (lower genomic coordinates), whereas positive values indicate downstream localization.

## Code Availability

Executable notebook code spanning all key analyses across main and supplementary figure panels is provided as a GitHub repository https://github.com/modiashini/HRDbackup

## Acknowledgements

We are grateful to Mehmet K. Samur and Alan D’Andrea for the insightful discussions and suggestions. A.Z. received funding from thePaula and Rodger Riney Foundation. A.Z. and G.P. received funding from NIH-NCI 5R01 CA262710-05 and 5P01CA155258-12

