## Supplementary materials for "Pan-Cancer Genomic Scars of Alternative End Joining and Single-Strand Annealing"

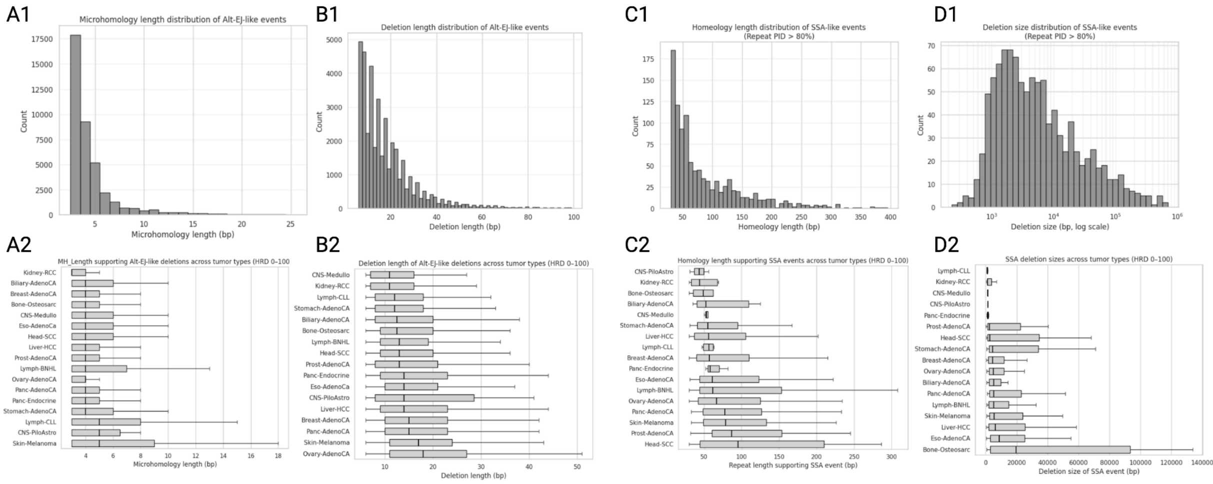

**Fig. S1 | Homology length and deletion size characteristics of Alt-EJ– and SSA-like events.**
**a,b,** Distribution of microhomology length **(a1)** and deletion length **(b1)** for Alt-EJ-like events across all tumors. **a2,b2,** Distribution of microhomology length **(a2)** and deletion length **(b2)** for Alt-EJ-like events for each tumor type. Box plots show median and interquartile range, with whiskers extending to 1.5× the interquartile range. **c,d,** Distribution of homeology length (repeat sequence identity >80%) **(c1)** and deletion size (log scale) **(d1)** for SSA-like events across all tumors. **c2,d2,** Distribution of homeology length **(c2)** and deletion size **(d2)** for SSA-like events for each tumor type.

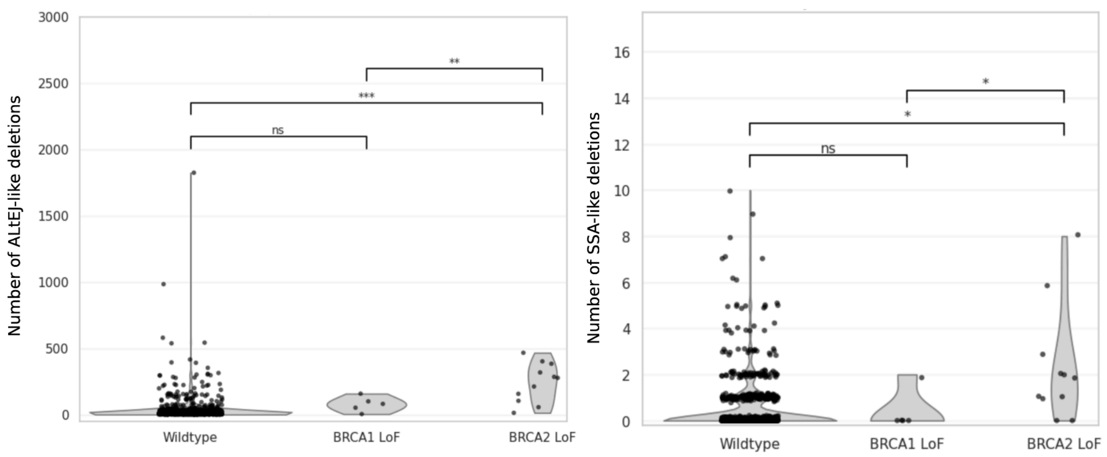

**Fig. S2 | Increased Alt-EJ and SSA usage in BRCA2 but not BRCA1-deficient tumors.**

**a,** Number of Alt-EJ-like deletions per tumor in wild-type, BRCA1 loss-of-function (LoF) and BRCA2 LoF groups. **b,** Number of SSA-like deletions per tumor across the same groups. Points represent individual tumors overlaid on violin plots showing the distribution within each group. Statistical significance between groups is indicated via Welch's t-tests (ns, not significant; *P < 0.05; **P < 0.01; ***P < 0.001). Variants were classified as LoF if they were frameshift, stop codon gain, splice variant, or start codon loss.

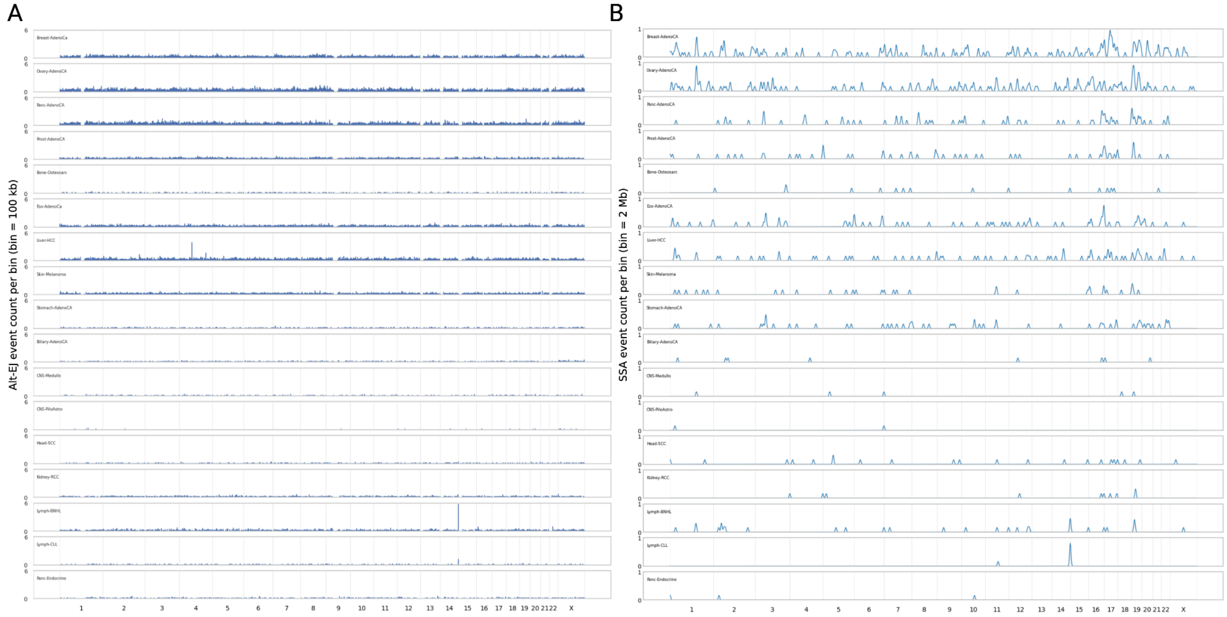

**Fig. S3 | Genome-wide distribution of Alt-EJ- and SSA-like deletions across tumor types.**
**a,** Genome-wide distribution of Alt-EJ-like deletions across individual tumor types, shown as event counts per 100 kb bin with chromosomes plotted sequentially along a continuous hg19 coordinate axis. Each row represents a tumor type. **b,** Genome-wide distribution of SSA-like deletions across the same tumor types, shown as event counts per 2 Mb bin. Each row represents a tumor type.

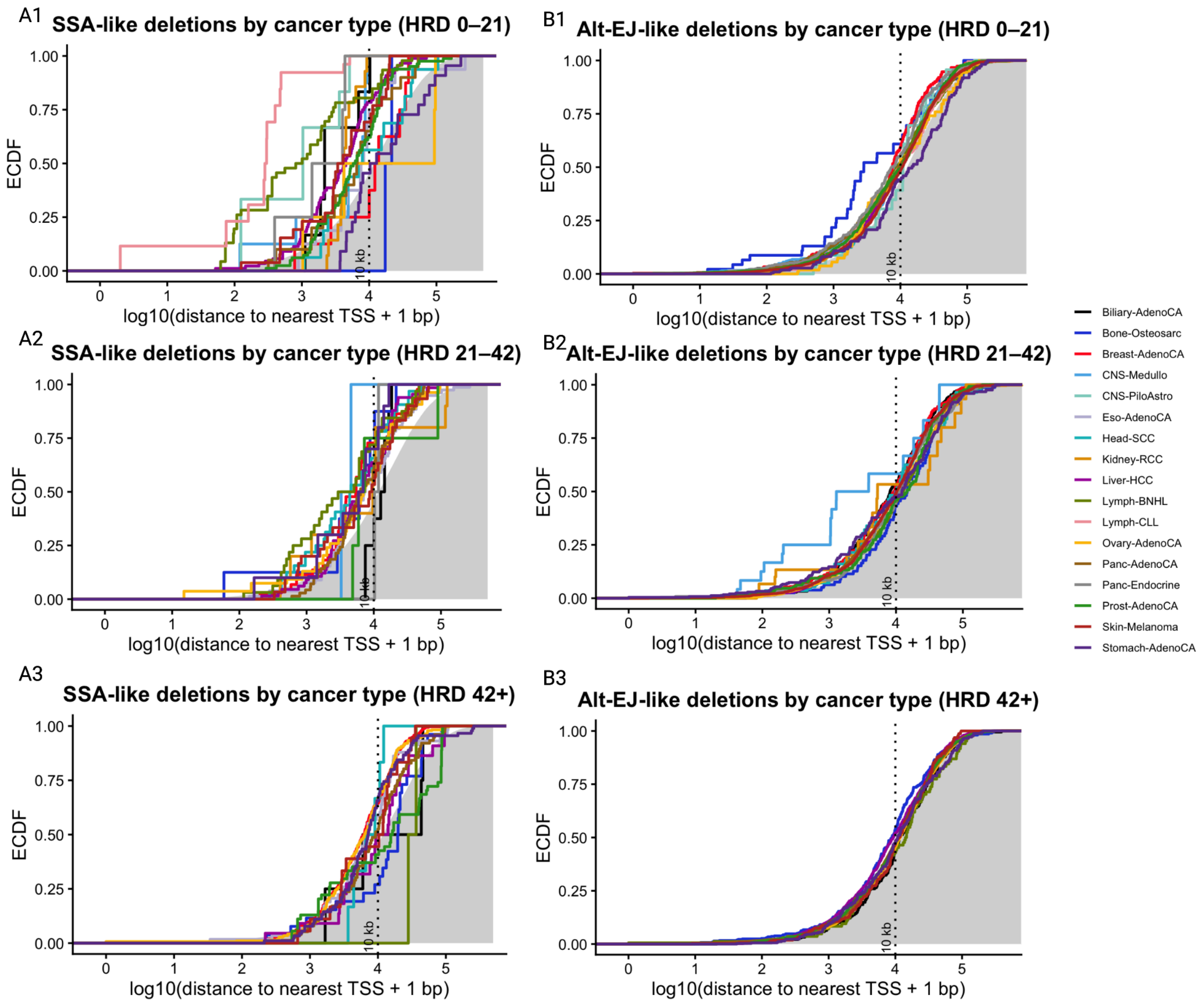

**Fig. S4 | Distribution of SSA- and Alt-EJ-like deletion breakpoints relative to transcription start sites (TSS).** **a1-a3**, Empirical cumulative distribution functions (ECDFs) of distances from SSA-like deletion breakpoints to the nearest TSS, shown for individual cancer types within **(a1)** HRD 0-21, **(a2)** HRD 21-42, and **(a3)** HRD >42 tumors. **b1-b3,** ECDFs of distances from Alt-EJ-like deletion breakpoints to the nearest TSS, shown for individual cancer types within **(b1)** HRD 0-21, **(b2)** HRD 21-42, and **(b3)** HRD >42 tumors. Distances are plotted as log10(distance to nearest TSS + 1 bp). Each colored line represents a cancer type. The vertical dashed line indicates 10 kb from the TSS. Grey shading denotes the ECDF of background deletions (deletions >100 bp for panels a1-a3; deletions <100 bp for panels b1-b3 aggregated across all tumors and HRD groups, providing a reference distribution for comparison.

| Cancer | HRD bin | Number of SSA events | Number of SSA events <250 bp from a TSS | Proportion of SSA events <250 bp from a TSS | Number of tumors with ≥1 SSA event |
|---|---|---|---|---|---|
| **Biliary-AdenoCA** | HRD 0-21 | 3 | 0 | 0.00 | 2 |
| **Bone-Osteosarc** | HRD 0-21 | 1 | 0 | 0.00 | 1 |
| **Breast-AdenoCA** | HRD 0-21 | 4 | 0 | 0.00 | 3 |
| **CNS-Medullo** | HRD 0-21 | 4 | 1 | 0.25 | 4 |
| **CNS-PiloAstro** | HRD 0-21 | 2 | 1 | 0.50 | 2 |
| **Eso-AdenoCA** | HRD 0-21 | 8 | 0 | 0.00 | 6 |
| **Head-SCC** | HRD 0-21 | 8 | 0 | 0.00 | 7 |
| **Kidney-RCC** | HRD 0-21 | 6 | 0 | 0.00 | 6 |
| **Liver-HCC** | HRD 0-21 | 41 | 3 | 0.07 | 31 |
| **Lymph-BNHL** | HRD 0-21 | 15 | 5 | 0.33 | 13 |
| **Lymph-CLL** | HRD 0-21 | 6 | 3 | 0.50 | 6 |
| **Ovary-AdenoCA** | HRD 0-21 | 2 | 0 | 0.00 | 2 |
| **Panc-AdenoCA** | HRD 0-21 | 10 | 0 | 0.00 | 10 |
| **Panc-Endocrine** | HRD 0-21 | 2 | 0 | 0.00 | 2 |
| **Prost-AdenoCA** | HRD 0-21 | 38 | 0 | 0.00 | 29 |
| **Skin-Melanoma** | HRD 0-21 | 12 | 1 | 0.08 | 10 |
| **Biliary-AdenoCA** | HRD 21-42 | 3 | 0 | 0.00 | 2 |
| **Bone-Osteosarc** | HRD 21-42 | 4 | 1 | 0.25 | 3 |
| **Breast-AdenoCA** | HRD 21-42 | 19 | 1 | 0.05 | 13 |
| **CNS-Medullo** | HRD 21-42 | 1 | 0 | 0.00 | 1 |
| **CNS-PiloAstro** | HRD 21-42 | 0 | 0 | 0.00 | 0 |
| **Eso-AdenoCA** | HRD 21-42 | 37 | 1 | 0.03 | 22 |
| **Head-SCC** | HRD 21-42 | 13 | 0 | 0.00 | 7 |
| **Kidney-RCC** | HRD 21-42 | 4 | 0 | 0.00 | 4 |
| **Liver-HCC** | HRD 21-42 | 30 | 1 | 0.03 | 22 |

| | | | | | |
|---|---|---|---|---|---|
| **Lymph-BNHL** | HRD 21-42 | 14 | 1 | 0.07 | 6 |
| **Lymph-CLL** | HRD 21-42 | 0 | 0 | 0.00 | 0 |
| **Ovary-AdenoCA** | HRD 21-42 | 24 | 1 | 0.04 | 14 |
| **Panc-AdenoCA** | HRD 21-42 | 34 | 0 | 0.00 | 25 |
| **Panc-Endocrine** | HRD 21-42 | 1 | 0 | 0.00 | 1 |
| **Prost-AdenoCA** | HRD 21-42 | 2 | 0 | 0.00 | 2 |
| **Skin-Melanoma** | HRD 21-42 | 12 | 0 | 0.00 | 9 |
| **Biliary-AdenoCA** | HRD > 42 | 2 | 0 | 0.00 | 1 |
| **Bone-Osteosarc** | HRD > 42 | 11 | 1 | 0.09 | 7 |
| **Breast-AdenoCA** | HRD > 42 | 170 | 7 | 0.04 | 59 |
| **CNS-Medullo** | HRD > 42 | 0 | 0 | 0.00 | 0 |
| **CNS-PiloAstro** | HRD > 42 | 0 | 0 | 0.00 | 0 |
| **Eso-AdenoCA** | HRD > 42 | 29 | 1 | 0.03 | 13 |
| **Head-SCC** | HRD > 42 | 1 | 0 | 0.00 | 1 |
| **Kidney-RCC** | HRD > 42 | 0 | 0 | 0.00 | 0 |
| **Liver-HCC** | HRD > 42 | 11 | 1 | 0.09 | 8 |
| **Lymph-BNHL** | HRD > 42 | 1 | 0 | 0.00 | 1 |
| **Lymph-CLL** | HRD > 42 | 0 | 0 | 0.00 | 0 |
| **Ovary-AdenoCA** | HRD > 42 | 132 | 2 | 0.02 | 43 |
| **Panc-AdenoCA** | HRD > 42 | 31 | 0 | 0.00 | 17 |
| **Panc-Endocrine** | HRD > 42 | 0 | 0 | 0.00 | 0 |
| **Prost-AdenoCA** | HRD > 42 | 18 | 0 | 0.00 | 8 |
| **Skin-Melanoma** | HRD > 42 | 9 | 0 | 0.00 | 7 |

**Table S1 | Frequency of SSA-like deletions proximal to transcription start sites across cancer types and HRD bin.**
Number and proportion of SSA-like deletion events occurring within 250 bp of the nearest transcription start site (TSS) are shown for each cancer type and stratified by homologous recombination deficiency (HRD) bin (0-21, 21-42, >42). For each group, we report the total number of SSA events, the number and proportion occurring within ±250 bp of a TSS, and the number of tumors harboring at least one SSA event. The ±250 bp window corresponds to the central TSS-proximal region highlighted in Fig. S4. Across tumor types, TSS-proximal SSA events are most enriched in select HR-proficient cohorts (HRD 0-21), including lymphoid malignancies (Lymph-BNHL and Lymph-CLL).
